## Supporting Information for "Periodontitis promotes bacterial extracellular vesicle-induced neuroinflammation in the brain and trigeminal ganglion"

**This PDF file includes:**

Figures S1 to S7

Tables S1

Supporting information references

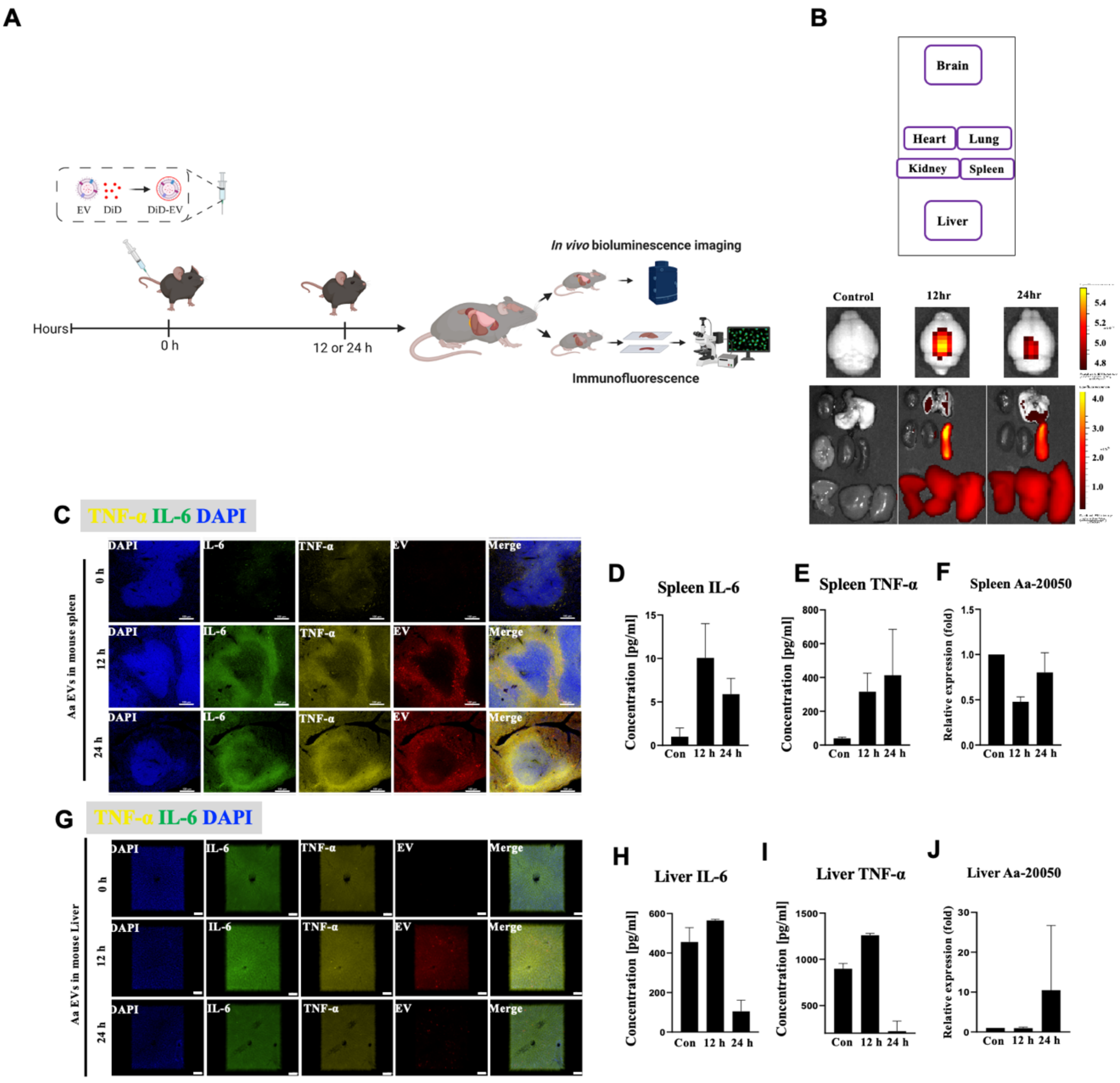


**S1 Fig. Aa EVs can spread to systemic organs following intravenous tail vein injection.** (A) General scheme of the in vivo experimental design. Purified Aa EVs were quantified using NTS device and calculated particle numbers were used for each injection. Six-week-old male mice were intravenously administered lipid tracer, DiD-labeled Aa EVs (approximately $4.5 \times 10^{10}$ particles) for the indicated period. (B) The brain, heart, lung, kidney, spleen, and liver were resected, and the distributions of DiD-labeled Aa EVs were determined using an in vivo imaging system (IVIS). Although weaker fluorescence intensity was observed in the brain and lungs than in the spleen and liver, the signals became strong after 12 to 24 h of i. v. injections; IVIS signals in the brain were measured separately. Previously, we showed that Aa EVs and exRNAs specifically induce IL-6 expression in microglial cells and TNF-α expression in the brain [1,2]. These proinflammatory cytokines were measured in the spleen and liver using immunofluorescence staining. Confocal image analysis was performed to detect IL-6 (green) and TNF-α (yellow) in spleen (C) and the liver (G). Red spots indicate Aa EVs. Spleen IL-6 (D), spleen TNF-α (E), liver IL-6 (H), and liver TNF-α (I) were quantified using ELISA. The levels of TNF-α and IL-6 were elevated in the spleen 12 and 24 h after injection, but not in the liver. To verify the successful Aa EV cargo delivery, one of the prominent miRNA-sized small RNAs present in Aa EV, Aa-20050 [3], was measured by qRT-PCR in the spleen (F) and liver (J). For qRT-PCR, Cel-miR-39-3p was added (spike-in) for normalization and found to accumulate in the liver after 24 h of injection but not in the spleen; this may be attributed to rapid degradation of small RNA in these organs due to its high RNase activity [4]. Data are presented as mean ± SD from five independent experiments. Scale bar = 100 μm. Depending on the organ, it may be possible for it to serve as both a target and an elimination organ [5]. Most Aa EVs accumulate in the spleen and liver but induce weaker proinflammatory cytokine release, suggesting that the liver and spleen might act as elimination organs for blood-borne bEVs and their cargo, including RNAs.

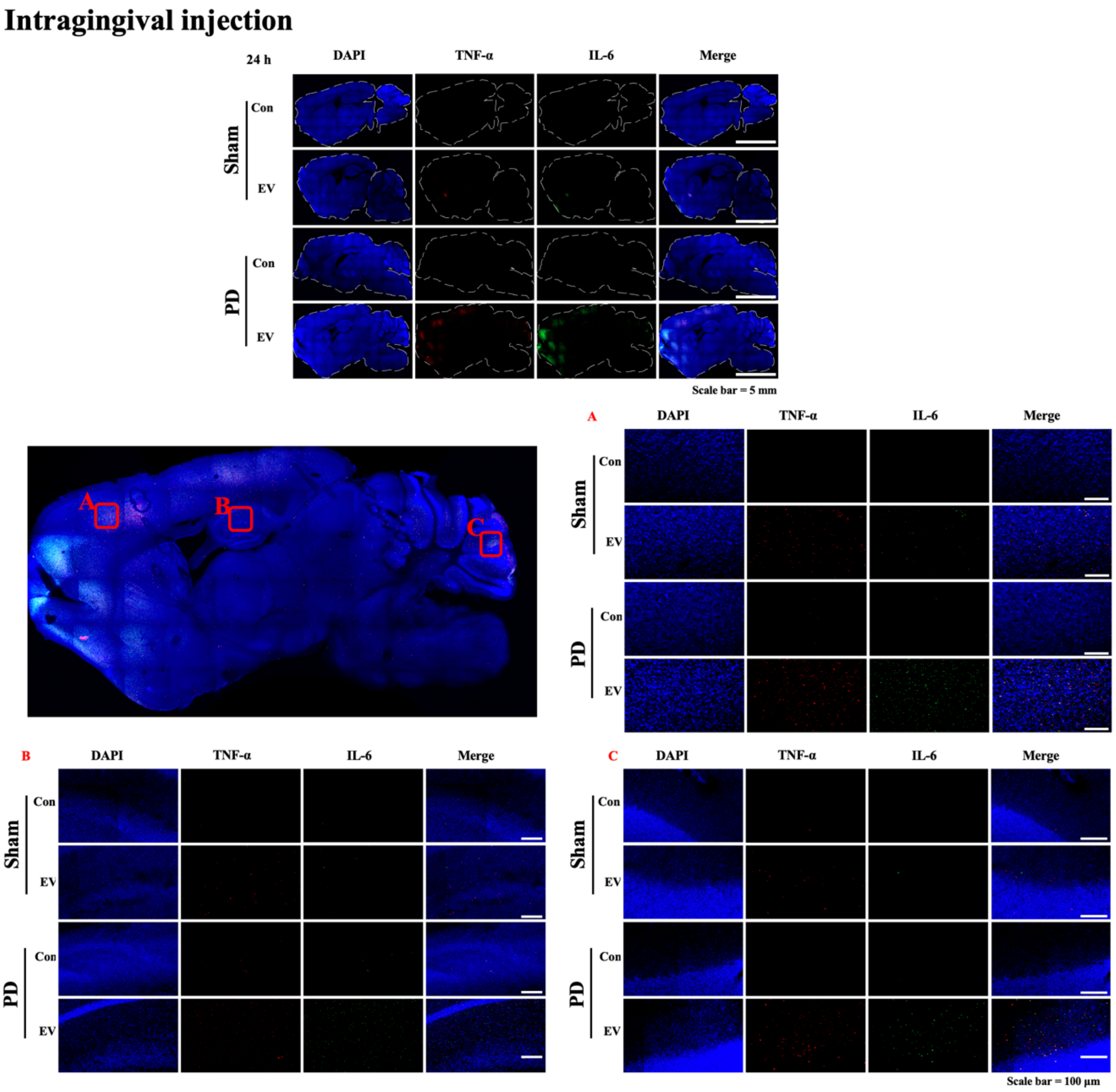


**S2 Fig. Aa EVs-induced proinflammatory cytokines secretion in PD model generated via ligature and subjected to intragingival injection.** DiD-labeled Aa EVs (approximately 4.5 × $10^{10}$ particles) were intragingivally injected to mice for 24 h. Confocal fluorescence image analysis showed increased Aa EV particles (red) in the brains of mice with ligature-induced PD. DAPI signals are shown in blue. Left panel is the sagittal section of the brain, showing a merged fluorescence image of Aa EV-injected PD mice. Boxes (A, cerebral cortex; B, hippocampus; C, cerebellum) in the middle image are magnified in the right panel.

### EV-soaked gel

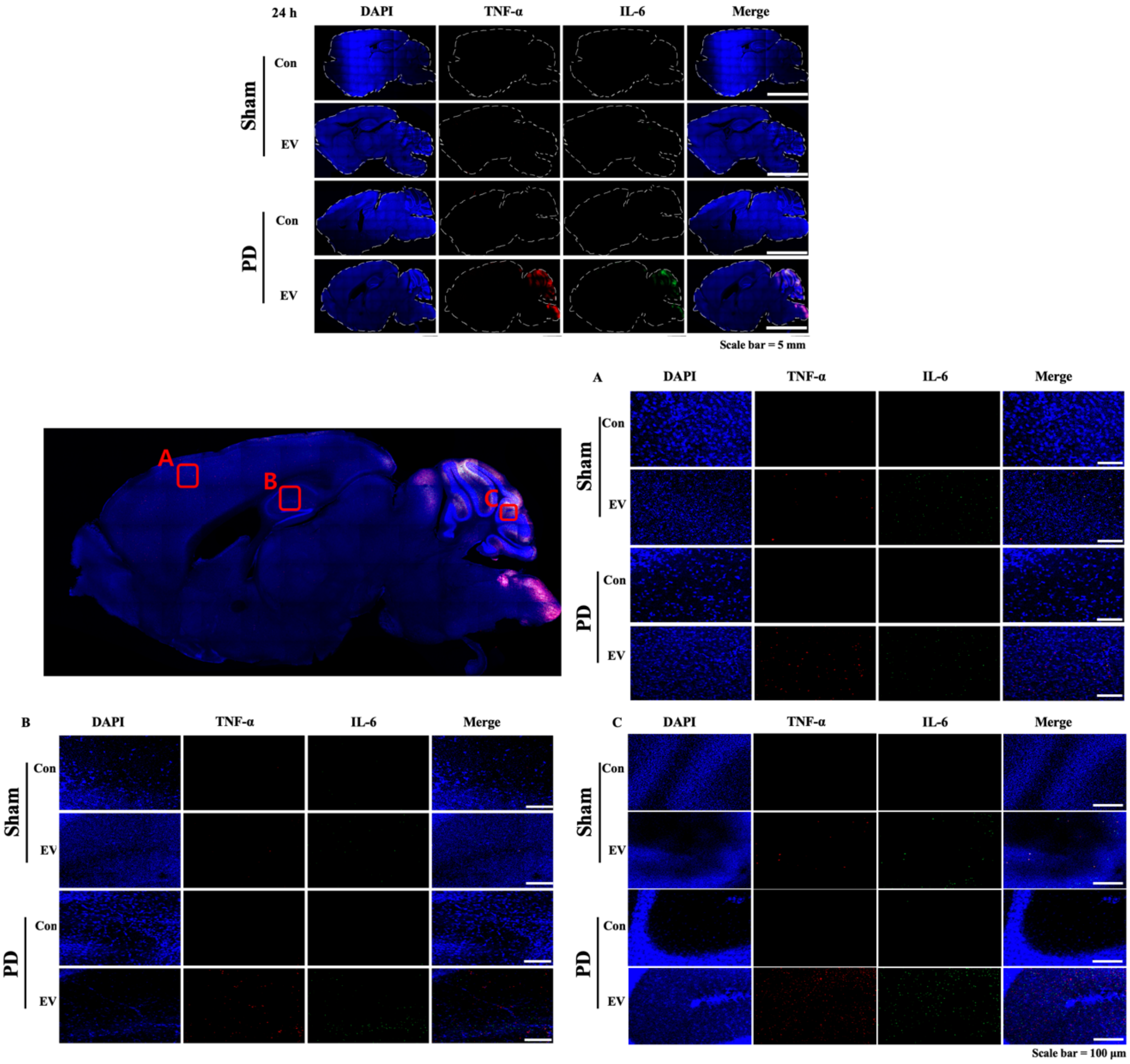


**S3 Fig. Aa EV-soaked gel-induced proinflammatory cytokine release in a mouse model of ligature-induced PD.** Gelatin gels including DiD-labeled Aa EVs (approximately 4.5 × $10^{10}$ particles) were administrated to mice for 24 h. Confocal fluorescence image analysis showed increased Aa EV particles (red) in the brains of mice with ligature-induced PD. DAPI signals are shown in blue. Left panel is the sagittal section of the brain, showing a merged fluorescence image of Aa EV-injected PD mice. Boxes (A, cerebral cortex; B, hippocampus; C, cerebellum) in the middle image are magnified in the right panel.

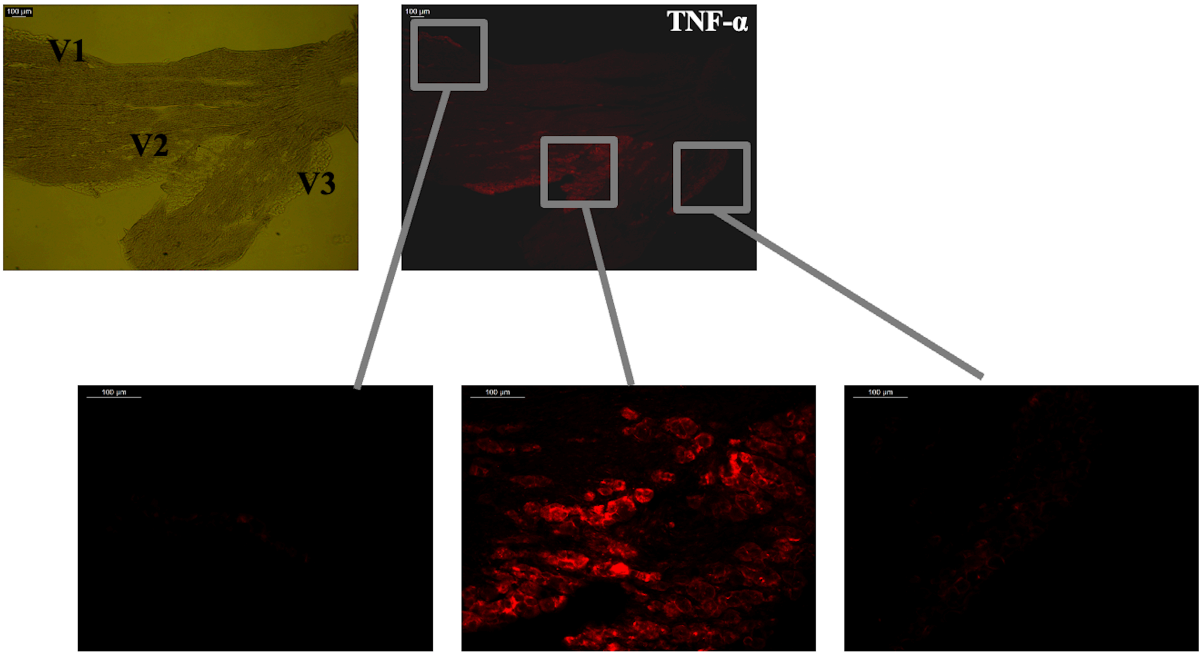


**S4 Fig. Aa EVs are directly transmitted by trigeminal ganglion (TG) neurons via axon terminal and stimulate the neurons.** The TG, which has three major branches: the ophthalmic (V1), maxillary (V2), and mandibular nerves (V3), relays painful sensation from the orofacial area and is unique among the somatosensory ganglia in terms of its topography, structure, composition, and possibly some functional properties of its cellular components. Epidermal injected Aa EVs were directly taken up by TG neurons and those Aa EVs activated TNF-α, specifically in the V2 region of TG. Boxes in the upper panel image are magnified in the lower panel. Scale bar = 100 µm.

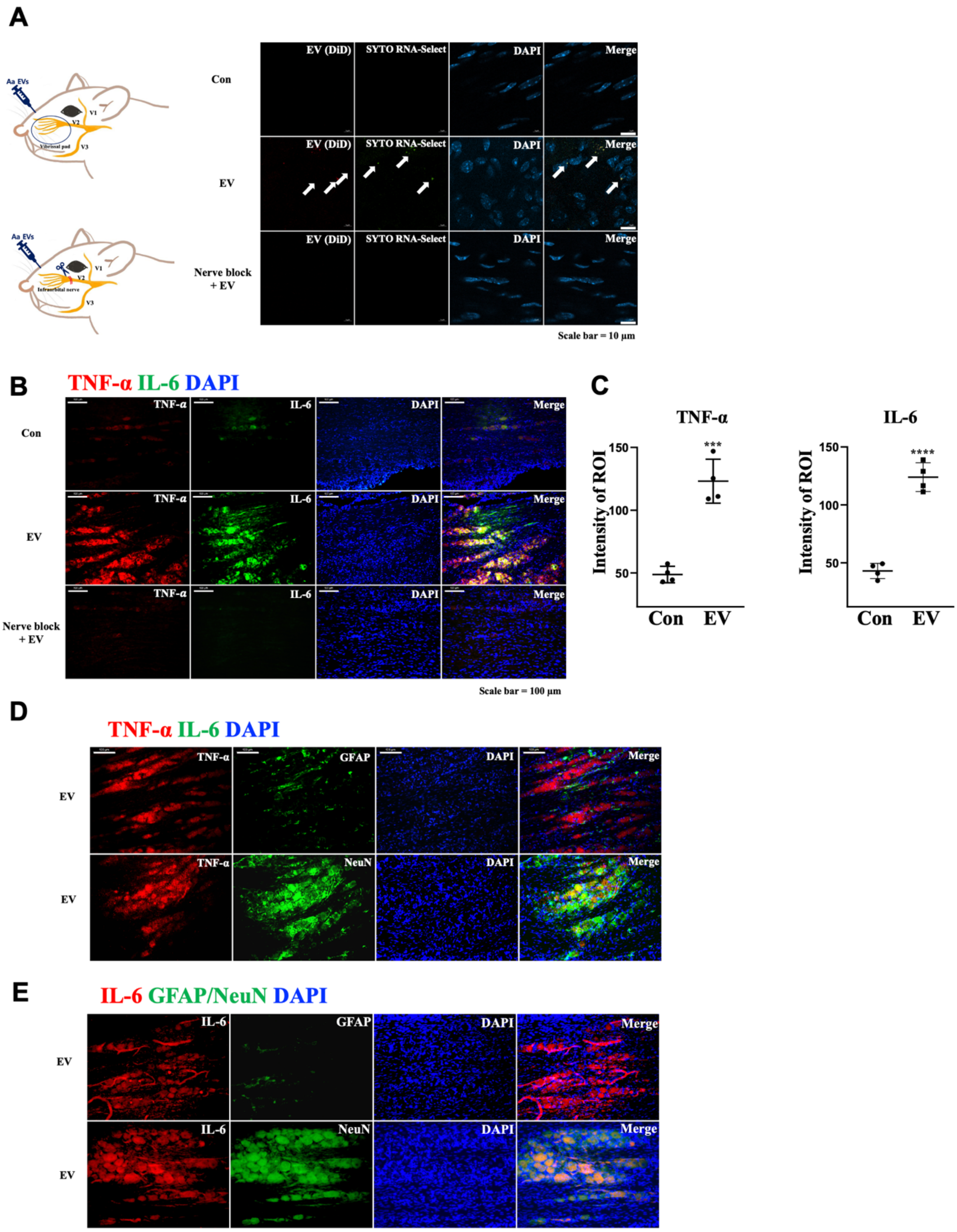


**S5 Fig. Aa EVs are directly transmitted by trigeminal ganglion (TG) neurons via axon terminal and stimulate the neurons.** (A) Pre-stained Aa EVs with lipid tracer dye DiD (red) and RNA-specific dye SYTO RNA-Select (green) were injected via direct infraorbital nerve (ION) injection. The V2 region of trigeminal ganglion, but not the nerve blocked TG, shows red spots (EV) and green spots (RNA inside EV) 24 h after injection (Scale bar = 10 μm). (B-C) Expression of proinflammatory cytokines TNF-α and IL-6 in the TG was compared with PBS control and ION-blocked TG. TNF-α and IL-6 positive cells were compared in Aa EVs administrated TG with PBS-control (Con) following ION injection (Scale bar = 100 μm). (D) TNF-α positive cells did not co-localize with astrocyte marker GFAP (green; upper panel), but did with neuronal marker NeuN (green; bottom panel) expressing cells following epidermal Aa EVs injection (Scale bar = 100 μm). (E) IL-6 (red) positive cells did not co-localize with astrocyte marker GFAP (green; upper panel), but did with neuronal marker NeuN (green; bottom panel) expressing cells following epidermal Aa EVs injection (Scale bar = 100 μm). The data represents four independent biological experiments and are presented as mean ± standard deviation (SD). ***$p \leq 0.001$, ****$p \leq 0.0001$ (Student's t-test).

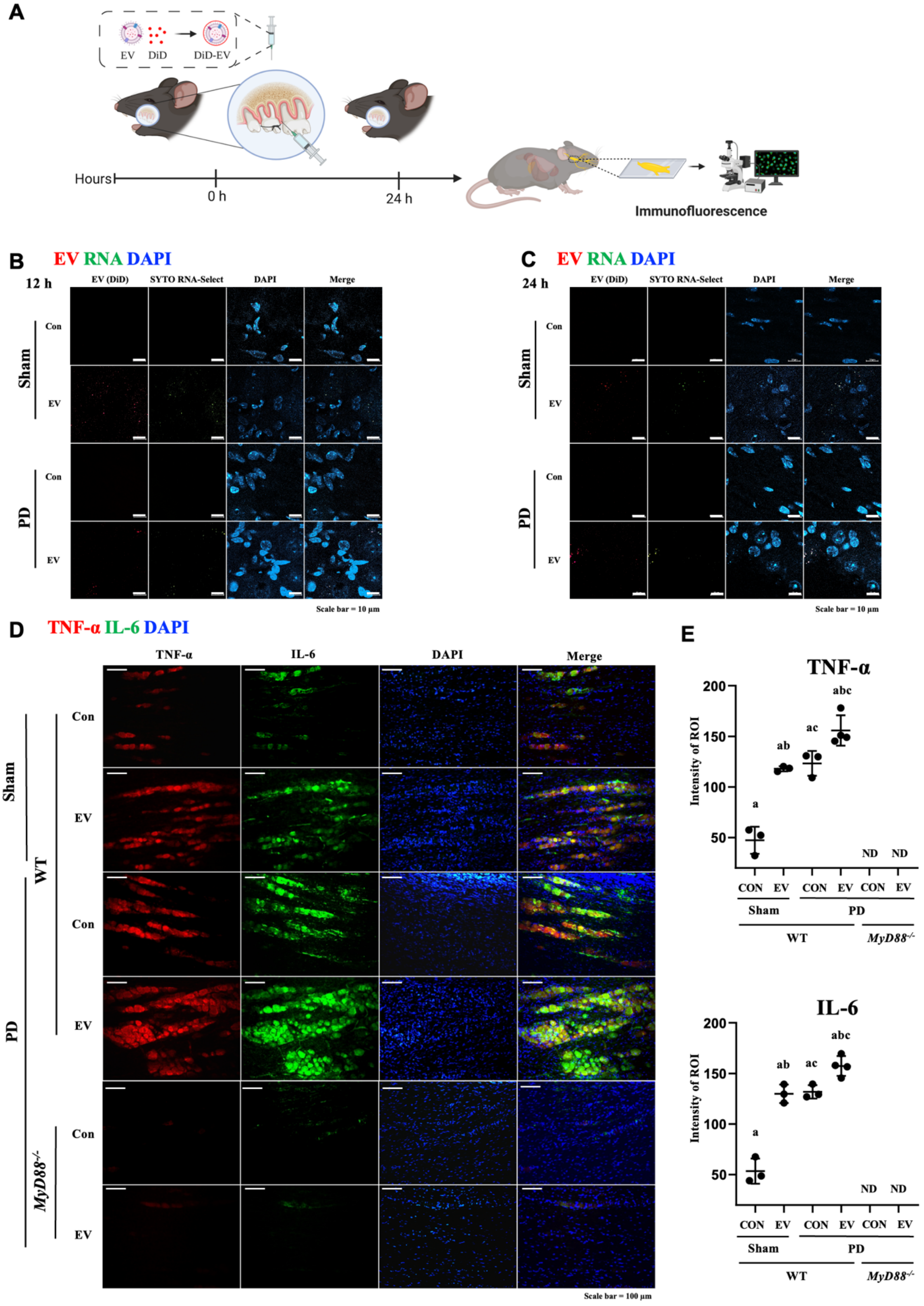


**S6 Fig. Aa EVs are directly transmitted by trigeminal ganglion (TG) neurons by gingival and Aa EV-induced proinflammatory cytokine expression is blocked in**

**TG of *MyD88$^{-/-}$* mice through intragingival administration.** DiD-labeled Aa EVs (approximately 4.5 × $10^{10}$ particles) were injected in WT and *MyD88$^{-/-}$* mice after ligature-induced necrosis. (A) General scheme of the in vivo experimental design. Pre-stained Aa EVs with lipid tracer dye DiD (red) and RNA-specific dye SYTO RNA-Select (green) were injected via intragingival injection. (B-C) The V2 region of trigeminal ganglion shows red spots (EV) and green spots (RNA inside EV) at 12 h (B) 24 h (C) after injection (scale bar = 10 μm). (D-E) Fluorescence microscopic image analysis revealed increased TNF-α (red) and IL-6 (green) level of TG in response to Aa EV intragingival injection, with a dramatic reduction of proinflammatory cytokines, TNF-α (red) and IL-6 (green) were observed by immunostaining in *MyD88$^{-/-}$* after 24 h of injection (Sham Con, n = 3; Sham EV, n = 3; PD Con, n = 3; PD EV, n = 4; *MyD88$^{-/-}$* Con, n = 3; *MyD88$^{-/-}$* EV n = 2). TNF-α and IL-6 positive cells were compared in Aa EV-administrated TG with PBS-control (Con) following intragingival injection (ND, non-detectable). The graphs are presented as mean ± standard deviation (SD). One-way ANOVA with Tukey's post-hoc test was used to compare each test group.

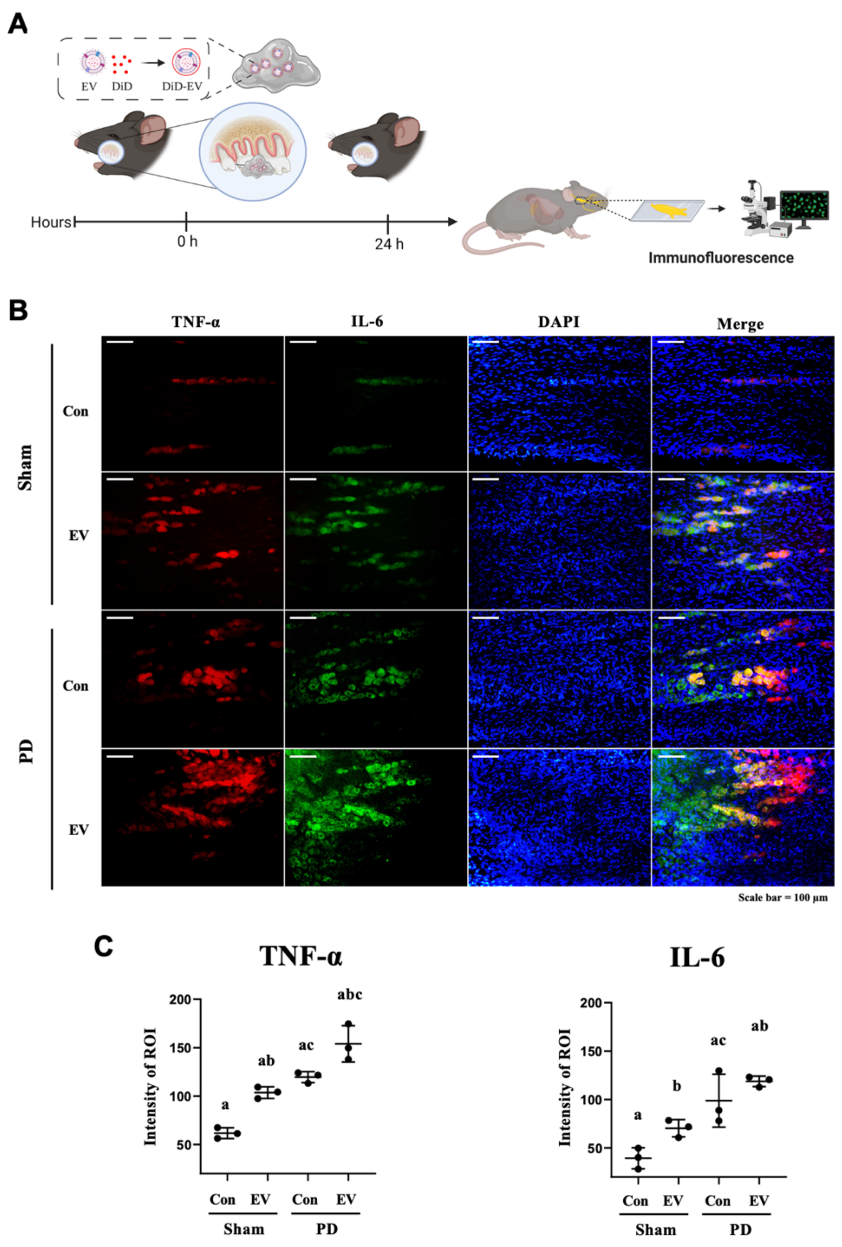


**S7 Fig. Aa EV-soaked gel induced proinflammatory cytokine release in trigeminal ganglion (TG) neurons of a mouse model of ligature-induced periodontal disease (PD).** (A) Schematic diagram of the in vivo experimental design. Gelatin gels including Aa EVs (approximately $4.5 \times 10^{10}$ particles) were administered to mice (Sham Con, n = 3; Sham EV, n = 3; PD Con, n = 3; PD EV, n = 3). (B) Immunostaining of TNF-α (red), IL-6 (green), and DAPI (blue) in the TG is shown. Scale bar = 100 μm. (C) TNF-α and IL-6 in TG quantification using ELISA showed their increased level in response to Aa EV-soaked gel. The graphs are presented as mean ± standard deviation (SD). One-way ANOVA with Tukey's post-hoc test was used to compare each test group.

**Table S1. Passive and active membrane properties of small-sized trigeminal ganglion (TG) neurons.**

| | $C_m$ (pF) | RMP (mV) | $R_{in}$ (MΩ) | AP amplitude (mV) | AP duration (ms) | AHP (mV) | $\tau$ of AHP (ms) | n |
|---|---|---|---|---|---|---|---|---|
| Control (PBS) | 14.9 ± 0.3 | -57.1 ± 1.5 | 691.9 ± 52.4 | 110.6 ± 2.0 | 4.6 ± 0.2 | 12.5 ± 0.9 | 54.5 ± 3.7 | 52 |
| Aa EV | 14.3 ± 0.3 | -57.4 ± 1.0 | 737.1 ± 56.4 | 111.3 ± 1.9 | 4.4 ± 0.3 | 11.6 ± 0.7 | 52.5 ± 6.2 | 50 |
| *p*-value[a] | 0.124 | 0.855 | 0.559 | 0.807 | 0.677 | 0.413 | 0.697 | |

Data represent the mean and SEM.

$C_m$; membrane capacitance, RMP; resting membrane potential, $R_{in}$: input resistance, AP: action potential, AHP: after hyperpolarization, τ: decay time constant

[a]unpaired t-test.

**Supporting information references**
